# On the accuracy of image registration in portable low-field 3D brain MRI

**DOI:** 10.64898/2026.02.11.705413

**Authors:** J. Eugenio Iglesias, Ian P. Johnson, Jonathan Williams-Ramirez, Dina Zemlyanker, Lin Tian, Karthik Gopinath, Mark Olchanyi, Ava D. Farnan, Amelia Demopoulos, Matthew S. Rosen, Kevin N. Sheth, Adam de Havenon, W. Taylor Kimberly, Annabel Sorby-Adams, Alzheimer’s Disease Neuroimaging Initiative (ADNI)

## Abstract

Portable low-field MRI offers an affordable and mobile alternative to conventional high-field scanners, enabling imaging in point-of-care and resource-limited settings. However, its lower signal-to-noise ratio, reduced resolution, and acquisition artifacts raise concerns about the accuracy of standard image registration methods. Reliable registration is critical for a wide range of emerging applications, including frequent brain monitoring, assessment of neurodegenerative disease progression, and evaluation of treatment effects such as those of Alzheimer’s therapeutics. In this work, we systematically evaluated state-of-the-art registration approaches on simulated low-field scans (obtained by downsampling high-field images) and on real low-field brain MRI data. We compared three representative approaches: classical optimization (NiftyReg), learning-based registration (SynthMorph), and synthesis-based registration (SynthSR+NiftyReg). Using downsampled high-field scans, all methods performed well, achieving high Dice scores and smooth deformation fields, indicating that reduced resolution alone does not hinder registration. In contrast, real low-field data exhibited lower accuracy, primarily due to geometric distortion and other acquisition-specific artifacts. Among the tested approaches, the synthesis-based pipeline achieved the most robust performance across subjects and modalities. Overall, existing algorithms can accommodate resolution limitations, however, future methods could further enhance coregistration by explicitly addressing the distortions present in low-field MRI scans.

## Introduction

Image registration seeks to determine a spatial transformation that aligns one image (the “moving” image) to another (the “fixed” image), thereby establishing pixel-(2D) or voxel-wise (3D) correspondence of their contents. In medical imaging, this foundational task underpins operations such as segmentation propagation, atlas-based quantification, longitudinal change detection, multimodal fusion, and planning of image-guided interventions^1–6^.

In human neuroimaging using brain magnetic resonance imaging (MRI), registration is ubiquitous. It enables spatial normalization of individual scans to a common template (such as MNI space^7^) for group analyses, voxel-based morphometry, surface-based analyses, and longitudinal within-subject alignment to assess subtle structural change over time (e.g., atrophy or ventricular enlargement)^8–10^. Rigid and affine registration are typically employed to correct for global positioning, while nonrigid (deformable) registration accommodates anatomical variability and local deformations, enabling fine-scale alignment across subjects and timepoints^11–13^.

For over two decades, brain MRI registration methods relied primarily on classical optimization-based frameworks: one formulates an energy minimization problem combining a similarity term (e.g., sum of squared differences, normalized cross-correlation, mutual information) and a regularizer (such as bending energy) characterizing the deformation field^14,15^. Popular tools include ANTs^13^, NiftyReg^16^, Elastix^17^, FNIRT^18^, or SPM^19^, which iteratively optimize transformation parameters for each image pair from scratch. These classical approaches offer well-understood behavior, explicit regularization control, and guarantee of invertibility or topology preservation (in diffeomorphic variants). These features have made such approaches gold standards in neuroimaging pipelines. For example, the UK BioBank, which is currently the largest initiative for population-based imaging of the human brain with multi-modal MRI, relies on FNIRT for atlas registration^20^.

In the last 10 years, registration has been transformed by deep learning techniques. These methods train convolutional neural networks (CNNs^21^) or, more recently, Transformers^22^ to predict deformation fields in one forward pass, thereby amortizing the computational cost of per-pair optimization. Early learning approaches were supervised, learning from synthetic deformation fields^23^ or from classical registration outputs^24^. Newer methods, on the other hand, adopt unsupervised or weakly supervised losses that mirror classical registrations (e.g., similarity + smoothness criteria) and allow training end-to-end^25–28^. Learning-based registration is orders of magnitude faster than classical methods at inference time, and many models incorporate innovations such as multi-scale (progressive) warping, segmentation loss during training, or uncertainty quantification^25,29,30^. Although numerous neuroimaging studies continue to rely on classical registration due to its mature theoretical guarantees (especially for diffeomorphic alignment), learning-based approaches are increasingly competitive in accuracy; see, e.g., TransMorph^31^ (based on transformers), Diffusemorph^32^ (based on diffusion models^33^), or uniGradICON^34^ (a foundation model for registration).

Crucially, both classical and learning-based registration methods perform extremely well on the standard 1 mm isotropic brain MRI scans typically used in research settings (e.g., high-field 1.5 T or 3 T acquisitions). In such high-quality data, metrics of alignment (e.g., Dice overlap of segmentations, target registration error) often exceed 0.9 and deformation fields remain well-behaved^9^. These performance levels support large-scale morphometric studies, longitudinal neurodegenerative tracking, and inter-subject atlas mapping.

In parallel, a new paradigm has emerged with the advent of portable, low-field MRI systems (e.g., <0.1 T), exemplified by Hyperfine’s 64 mT Swoop^®^ scanner. These devices offer substantial advantages: lower cost, reduced infrastructure requirements (no superconducting magnet or heavy shielding), low power consumption, and compact, mobile designs that enable point-of-care imaging. Early clinical applications include bedside neuroimaging in intensive-care settings and deployment in low-resource or rural environments where conventional MRI is unavailable^35–37^. By extending access beyond traditional hospital suites, portable MRI has the potential to democratize neuroimaging and support broader global-health and community-based applications.

Nevertheless, the low magnetic field regime brings significant challenges. Reduced signal-to-noise ratio, lower spatial resolution, relaxed gradient hardware, partial Fourier, and accelerated acquisition trade-offs all contribute towards degrading image quality. Moreover, portable scanners frequently exhibit increased geometric distortion, gradient non-linearity, radio-frequency coil inhomogeneity, and susceptibility to electromagnetic interference due to minimal shielding or portable designs^38^. These acquisition-specific artifacts compromise the spatial fidelity of the imaging volume and pose new obstacles for registration, especially in longitudinal contexts where precise correspondence across time is essential.

Accurate image registration (both cross-sectional and longitudinal) is fundamental to unlocking the full potential of portable, low-field brain MRI, both on the affordability and point-of-care imaging fronts. From the affordability perspective, registration enables reliable measurement of subtle structural changes across time in large-scale or frequent imaging scenarios – for example, monitoring disease progression or therapeutic response in Alzheimer’s trials, where conventional high-field MRI is costly or logistically constrained. Registration also enables standardized population-level analyses by aligning low-field data from different subjects and sites, facilitating epidemiological and preventive screening studies. On the portability front, accurate registration is critical for tracking recovery trajectories in patients imaged at the bedside after stroke, traumatic brain injury, or neurosurgical intervention, where scans may differ substantially in positioning and geometry. Similarly, in emergency department or low-resource settings, registration can assist in aligning multimodal or serial scans to detect evolving pathology or assess treatment effects. Despite this broad potential, the accuracy of registration in low-field MRI remains poorly characterized, and it is unclear whether current methods achieve the precision required for these applications.

In this work, we systematically investigate the problem of image registration in portable, low-field brain MRI. We present an experimental framework to evaluate three representative state-of-the-art registration paradigms: classical optimization-based, learning-based, and synthesis-based approaches. Evaluation relies on both simulated low-resolution data derived from high-field MRI and real low-field acquisitions, including longitudinal scans. This framework allows us to disentangle the effects of resolution, noise, and acquisition-specific artifacts on registration accuracy and deformation regularity. Beyond benchmarking, we propose practical strategies for achieving robust alignment in low-field settings, including synthesis-based and longitudinal registration techniques. By delineating the capabilities and limitations of current methods, this work provides a foundation for developing registration algorithms tailored to portable MRI and for designing future studies that leverage low-field imaging for accessible, quantitative neuroimaging.

## Results

### Effect of resolution

We first evaluate the intrinsic capability of existing registration methods to align anatomical structures at the low resolutions achievable with portable low-field devices, independently of other artifacts specific to such scanners. To this end, we artificially downsampled ∼1 mm isotropic T1 and FLAIR scans from 40 unique subject pairs in the ADNI3 dataset (which yields images with almost no noise due to the antialiasing filter) and registered them using several methods. The resulting deformation fields were then interpolated back to the original 1 mm isotropic resolution of the native scans. These deformation fields were applied to warp the labels of the moving images, which were subsequently compared to the labels of the fixed images using Dice similarity coefficients.

We compare registration performance at two different resolutions: 1.6 × 1.6 × 5.0 mm axial, which corresponds to the default resolution of the “stock” sequences distributed with the Hyperfine Swoop^®^ scanner, and 3 mm isotropic, which represents the highest isotropic resolution that was achievable within 3–4 minutes of scanning at the time of this study, and which has been shown to yield good segmentation performance^39^. We evaluate two modality scenarios: intra-modality (T1 to T1) and inter-modality (T1 to FLAIR). In terms of registration methods, we consider: *(i)* a classical iterative approach (NiftyReg^16^), using LNCC^13^ as the similarity function for intra-modality registration and mutual information^40^ for inter-modality; *(ii)* a modern learning-based approach (SynthMorph^41^), which is both modality- and resolution-agnostic; and *(iii)* a synthesis-based approach combining SynthSR^42^ (which predicts 1 mm isotropic T1 images from scans of arbitrary contrast and resolution) with NiftyReg using LNCC. Additionally, we report results obtained with the original 1 mm isotropic scans as a reference for the accuracy achievable with high-resolution data. These scans were registered directly with NiftyReg in the intra-modality scenario, or after SynthSR-based contrast harmonization in the inter-modality scenario.

We note that, in principle, one could use the deformations fields derived from the high-field scans as gold standard to evaluate the fields from the low-field data. However, the results obtained with this approach are unreliable because the field deformations are highly undefined in flat image regions (in the classical optical flow literature, this is know as the “aperture problem”^43^). Instead, we use a combination of Dice scores (measuring label overlap) and membrane energy (measuring the regularity of the deformation); for a given Dice score, lower membrane energies are preferred.

Figure 1 shows boxplots of the resulting Dice scores together with the membrane energies. In the intra-modality (T1–T1) scenario, registration accuracy remains high (median Dice above 0.8) even when directly registering the low-resolution images with NiftyReg. Performance is almost identical at 3 mm isotropic and at the stock resolution. Notably, the learning- and synthesis-based methods substantially close the gap with the high-resolution reference: SynthMorph matches the reference performance entirely, while the SynthSR/NiftyReg combination falls within just 1–2 Dice points. The membrane energies are slightly higher for optimization-based methods at higher resolutions (particularly for the high-resolution reference), but all remain within approximately 10% of each other.

**Figure 1.**
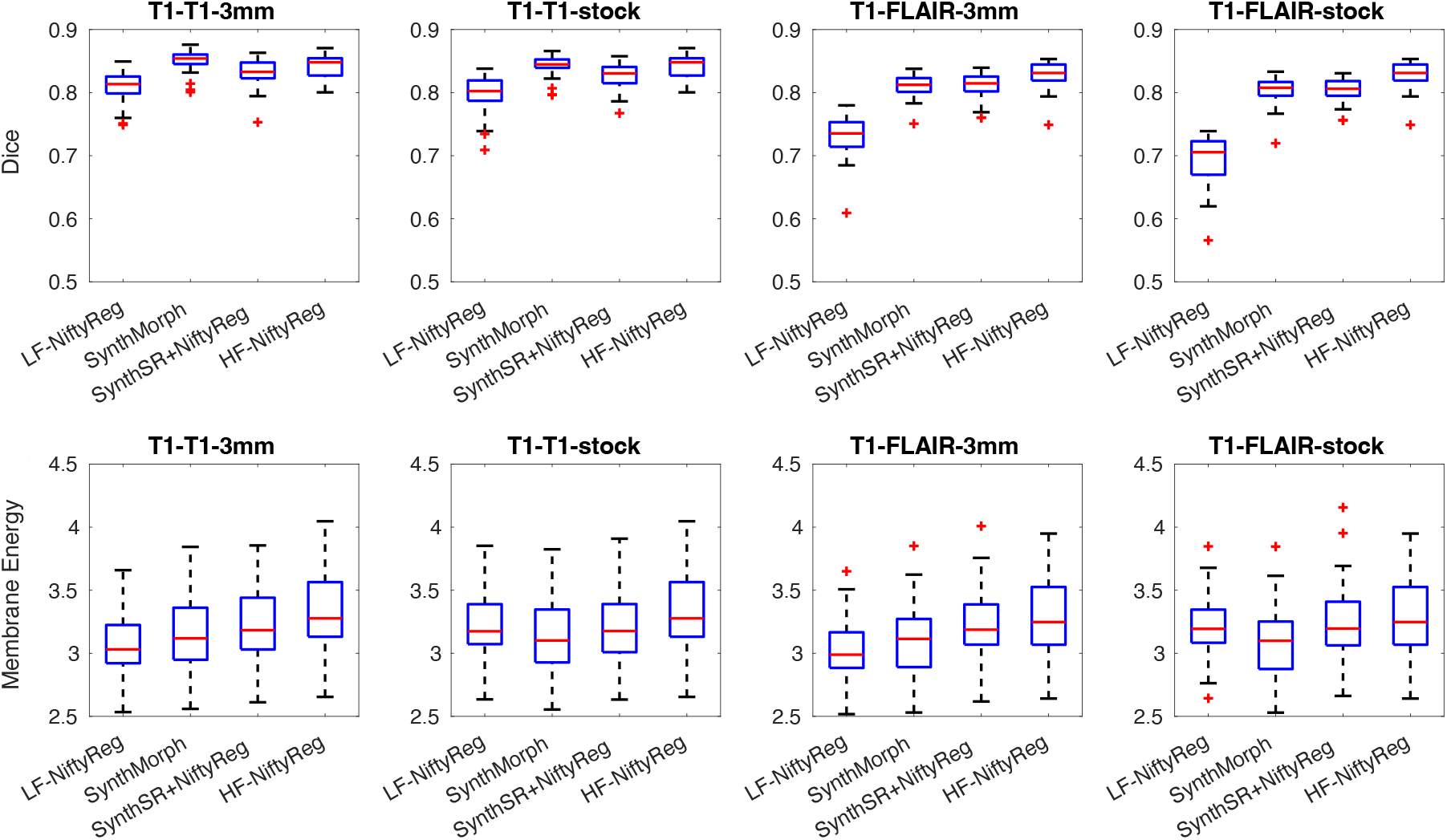
Box plots for Dice scores and membrane energy on downsampled ADNI dataset. We show results for intra- and inter-modality scenarios (T1-T1 and T1-FLAIR), at stock (1.6 × 1.6 × 5.0 mm axial) and 3 mm isotropic resolution. We compare direct registration of low-resolution data with NiftyReg (“LF-NiftyReg”) or SynthMorph; registration with NiftyReg of synthetic 1 mm T1 images obtained with SynthSR; and registration of the original 1 mm scans with NiftyReg (“HF-NiftyReg”). The boxes show medians (in red), interquartile ranges (IQR, in blue), whiskers (±1.5×IQR, in black), and outliers (red crosses).

In the inter-modality (T1–FLAIR) scenario, the behavior differs markedly. Classical optimization with NiftyReg performs poorly across modalities (a well-known limitation^44^) and this inaccuracy worsens at low resolutions, especially for anisotropic data from the stock sequence. In contrast, the learning- and synthesis-based approaches effectively handle the contrast differences, achieving accuracies comparable to those in the intra-modality setting. Remarkably, the high-resolution reference achieves nearly the same median Dice as the intra-modality case (83.1 vs. 84.8), despite the intermediate SynthSR synthesis step. Overall, SynthMorph and SynthSR/NiftyReg both yield median Dice scores above 0.8, reflecting robust registration performance in absolute terms, while maintaining membrane energies similar to those observed in the T1–T1 experiments.

Overall, reducing the resolution leads to only a modest loss in accuracy (∼ 2–3 Dice points relative to the high-resolution reference) and is accompanied by slightly lower membrane energies. This indicates that registration remains stable and anatomically consistent even at the resolutions attainable with portable low-field scanners. The broader impact of low-field imaging, including scanner-specific artifacts, is examined in the following sections.

### Effect of low-field scanning

Next, we evaluated the registration accuracy on real low-field data, using paired high- and low-field scans from the 40 cases in the Low-Field Outpatient Dataset, which comprises subjects with a clinical diagnosis of neurodegenerative disease that were recruited from an ambulatory outpatient neurology clinic. Each low-field scan was registered to a randomly selected different subject within the same pool, ensuring no repetitions and using the same registration methods and evaluation metrics as in the previous experiment.

A key difference from the simulated-resolution experiment is that the high- and low-field scans are not inherently aligned. Moreover, rigid registration cannot correct the nonlinear geometric distortions present in low-field acquisitions. This is illustrated with a phantom in Figure 2a–c: the apparent diameter at low field can be more than 5 mm off in some slices. While deformable registration can model these distortions, it requires an appropriate balance between transformation flexibility and image similarity. To identify suitable parameters, we first conducted a phantom experiment in which T1-, T2-, and FLAIR-weighted scans acquired at high field (3 T) were registered to corresponding low-field scans obtained across five sessions using different hardware and software versions of the portable scanner. Nonlinear registrations were performed in NiftyReg with LNCC, varying the B-spline control-point spacing between 5, 10, 15, 20, and 25 mm. We then plotted the final LNCC values and mean absolute log-Jacobian determinants as a function of spacing (Figure S1); we note that we use Jacobians instead of membrane energies because this should be a volume-preserving setting (since it is exactly the same scanned object). Based on these results, we selected a spacing of 20 mm as a compromise, since smaller spacings led to rapidly increasing irregularity. Qualitatively, the 20 mm transform provided sufficient flexibility to achieve accurate visual alignment (Figure 2d), and higher LNCC values seem to indicate overfitting.

**Figure 2.**
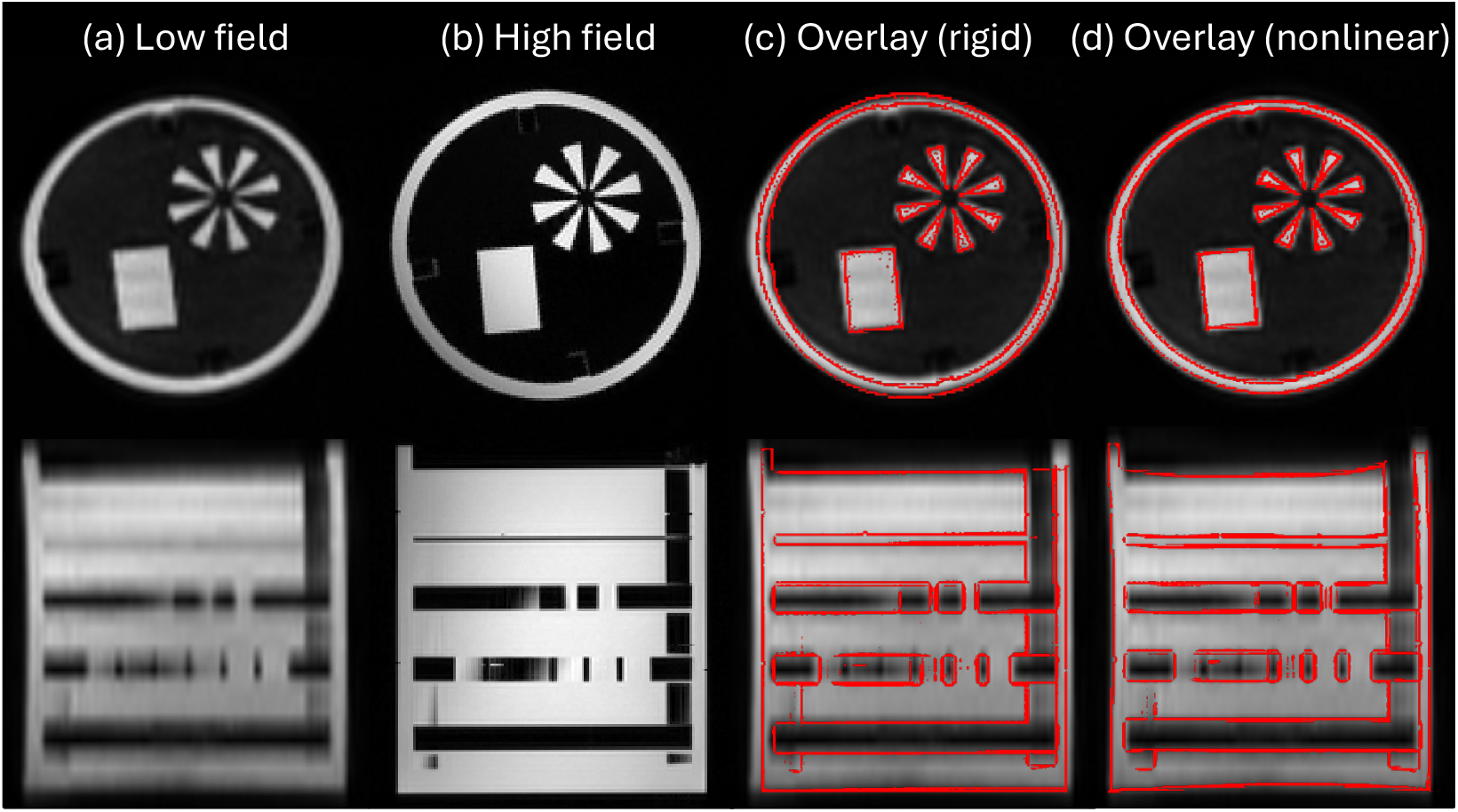
Orthogonal slices of a phantom scanned at low (a) and high field (b) using a T2-weighted sequence. Despite rigid registration, nonlinear misalignment remains due to geometric distortion at low field. This is illustrated by overlaying the edges of the high-field scan on top of the low-field scan (c); the error in the diameter of the phantom is as large as 5.3 mm (>4%). Nonrigid registration with a highly constrained deformation model (B-spline with 20 mm control spacing) largely mitigates this issue, without overfitting the deformation.

In light of the results from the phantom experiment, we used NiftyReg with LNCC and a B-spline deformation model with a 20 mm control-point spacing to derive ground-truth segmentations for the low-field data. Specifically, we registered each high-field scan to its corresponding low-field image using these settings and then warped the associated anatomical labels according to the resulting deformation fields. Although this approach introduces a small downward bias in Dice scores due to high- to low-field registration, it nevertheless provides the most reliable and consistent ground-truth approximation available for this dataset.

Figure 3 presents box plots of Dice scores and membrane energies. Compared with the downsampled experiment (which isolated the effect of reduced resolution), real low-field data show larger decreases in Dice and markedly higher membrane energies. This reflects the presence of characteristic low-field artifacts such as intensity inhomogeneities and geometric distortions, which produce more complex and irregular deformation fields. These effects are evident across all methods, whether registration is performed directly on low-field images (LF-NiftyReg and SynthMorph) or indirectly via SynthSR. In the latter case, the increased membrane energies likely stem from the fact that SynthSR must synthesize from geometrically distorted and noisy inputs, leading to less spatially smooth images than in the purely downsampled setup.

**Figure 3.**
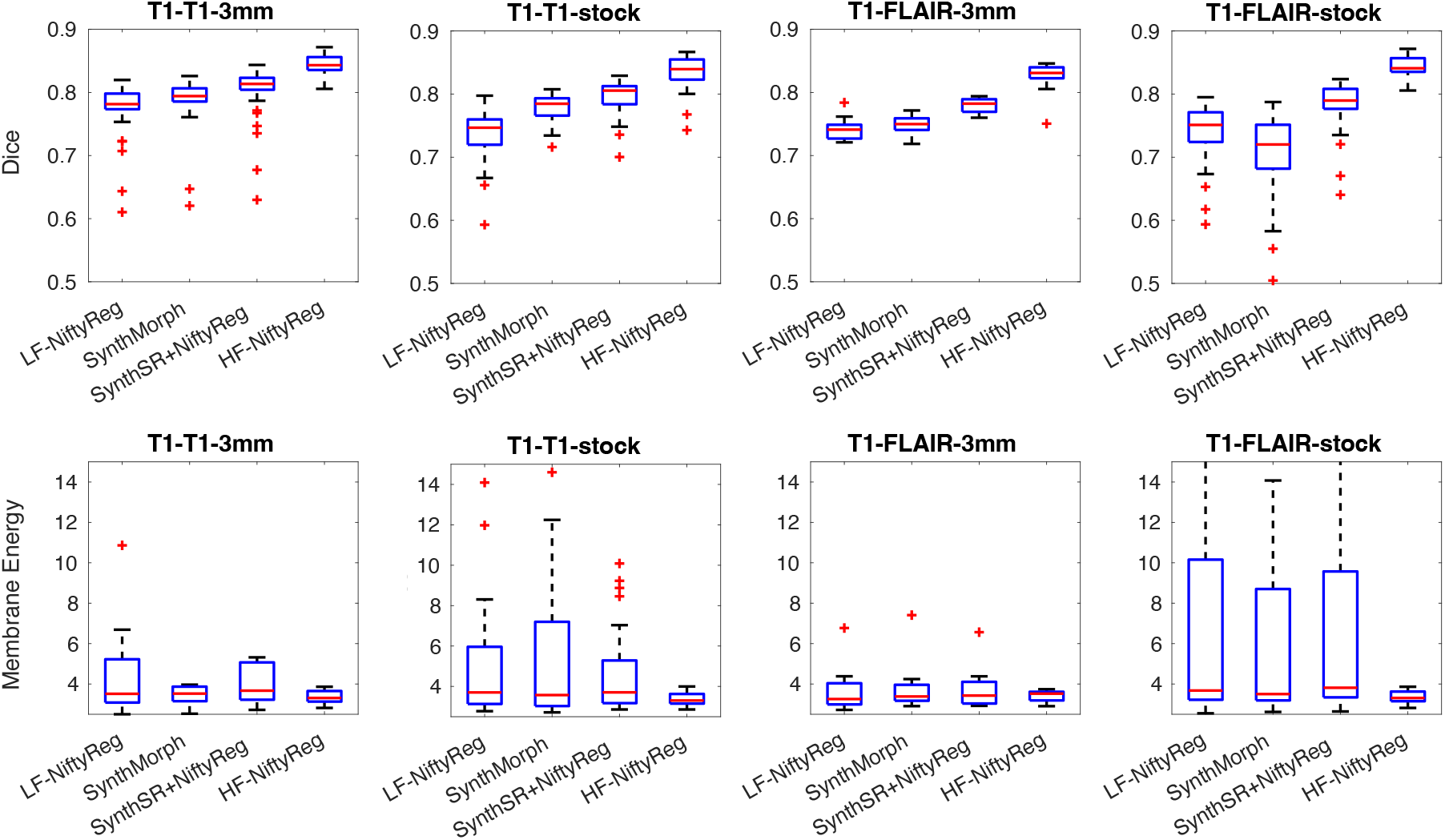
Box plots for Dice scores and membrane energy on Low-Field Outpatient Dataset. As in Figure 1, we show results for intra- and inter-modality scenarios (T1-T1 and T1-FLAIR), at stock (1.6 × 1.6 × 5.0 mm axial) and 3 mm isotropic resolution. We compare direct registration of low-resolution data with NiftyReg (“LF-NiftyReg”) or SynthMorph; registration with NiftyReg of synthetic 1 mm T1 images obtained with SynthSR; and registration of the original 1 mm scans with NiftyReg (“HF-NiftyReg”). The boxes show medians (in red), interquartile ranges (IQR, in blue), whiskers (± 1.5 × IQR, in black), and outliers (red crosses).

In the intra-modality (T1–T1) scenario, registration performance remains acceptable at 3 mm isotropic resolution, with median Dice scores close to 0.8 across methods (noting the small negative bias due to misregistration mentioned above). However, accuracy decreases notably when using the stock-resolution sequences. While consistent with previous observations on real data^39^, this trend was not observed with the simulated low-resolution data. This suggests that once geometric distortion and noise are introduced, the alignment task becomes substantially more challenging. Specifically, LF-NiftyReg drops to a median Dice of approximately 0.75, SynthMorph to 0.79, and SynthSR+NiftyReg marginally maintains values above 0.80.

A similar trend is observed in the inter-modality (T1–FLAIR) scenario. Classical optimization with LF-NiftyReg achieves ≈ 0.75 median Dice, while SynthMorph achieves 0.75 at 3 mm isotropic and 0.72 at the stock resolution. The SynthSR+NiftyReg pipeline remains the most robust, yielding median scores around 0.78–0.79. It should be noted that SynthMorph was not explicitly trained for low-field conditions and may not fully model the levels of noise and inhomogeneity present in these data. Overall, across both modality settings, LF-NiftyReg consistently performs worst, SynthMorph offers moderate improvement, and SynthSR+NiftyReg delivers the most stable results, maintaining median Dice values near 0.8 despite the challenges introduced by real low-field imaging. While these accuracies demonstrate that reliable registration is feasible at low field, they may still be insufficient for applications requiring very accurate voxel-level correspondence, such as longitudinal registration, which we assess in the following section.

### Registration of longitudinal scans at low-field

To evaluate registration accuracy in a longitudinal setting, we used scans from eight subjects who underwent both low- and high-field imaging sessions separated by 5.8 ± 2.8 months (see “Longitudinal Dataset” in the Methods). The low-field acquisitions included T1-weighted scans at two resolutions: 3 mm isotropic and the stock axial resolution (1.6 × 1.6 × 5.0 mm axial). As in the previous experiment, we applied a highly constrained nonlinear registration (B-spline with 20 mm control-point spacing) to transfer the gold-standard labels from the high- to the low-field scans. In addition, the high-field images were artificially downsampled to match the low-field resolutions, allowing us to disentangle the effects of resolution from those of real-world low-field artifacts.

Box plots of the Dice scores and membrane energies are shown in Figure 4. Because these are longitudinal scans of the same subjects, the reference Dice scores obtained from the high-field data are substantially higher than in the previous experiment (median: 0.92). As observed in the resolution experiments, the downsampled high-field images yield Dice values very close to this reference, with all three methods (LF-NiftyReg, SynthMorph, and SynthSR+NiftyReg) achieving median scores above 0.90. However, when real low-field scans are used, performance decreases noticeably, particularly at the anisotropic stock resolution. Among the evaluated methods, the SynthSR+NiftyReg combination performs best, reaching median Dice scores of 0.88–0.89. Nevertheless, even this approach produces individual cases with Dice scores that may be considered too low for longitudinal studies, where accurate registration between time points is critical for detecting subtle anatomical changes.

**Figure 4.**
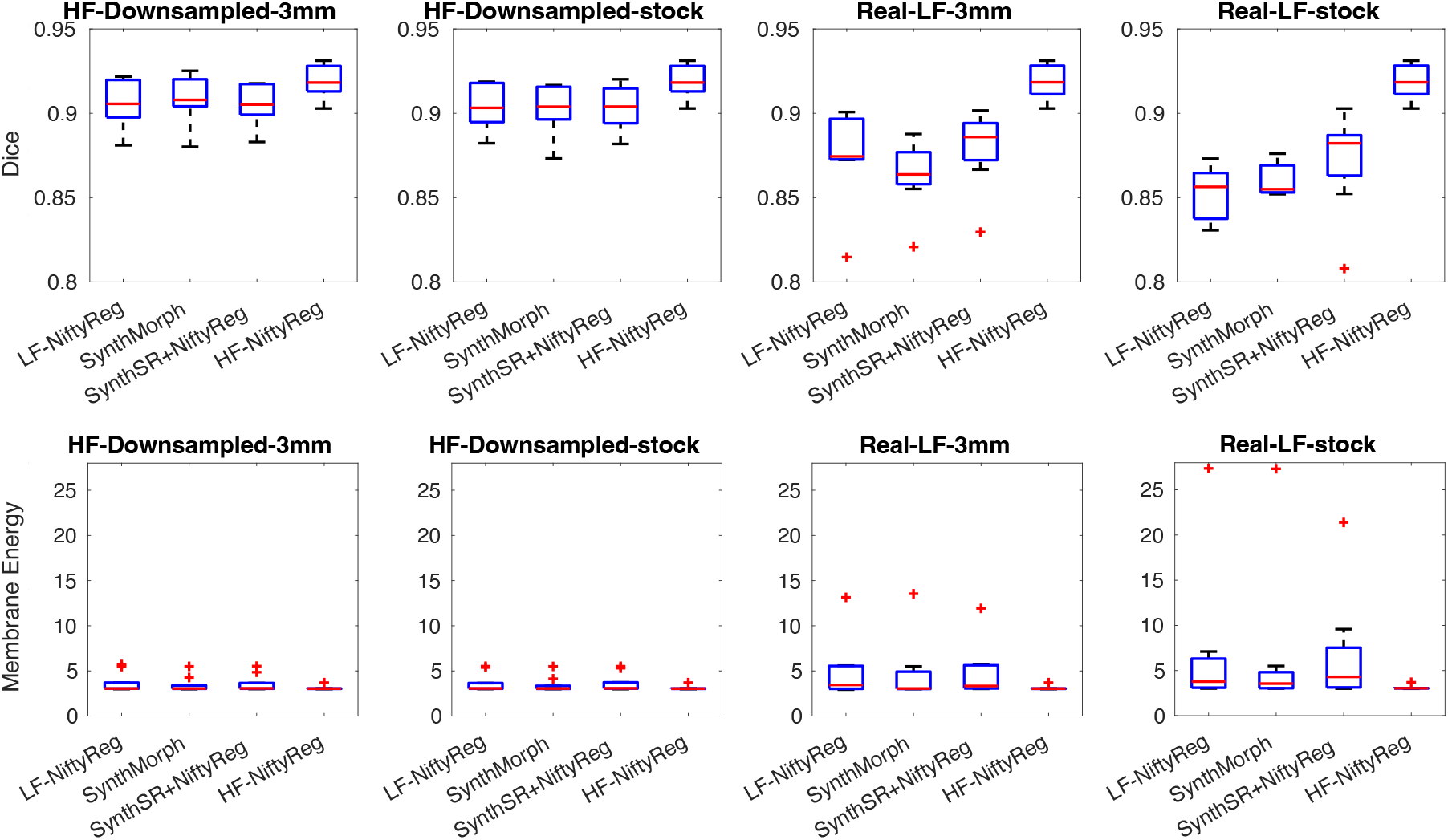
Box plots for Dice scores and membrane energy on longitudinal dataset. We show results at stock (1.6 × 1.6 × 5.0 mm axial) and 3 mm isotropic resolution, for downsampled and real low-field scans. As in previous experiments, we compare direct registration of low-resolution data with NiftyReg (“LF-NiftyReg”) or SynthMorph; registration with NiftyReg of synthetic 1 mm T1 images obtained with SynthSR; and registration of the original high-field with NiftyReg (“HF-NiftyReg”). The boxes show medians (in red), interquartile ranges (IQR, in blue), whiskers (±1.5×IQR, in black), and outliers (red crosses).

To further evaluate longitudinal registration performance, we tested it in a practical application scenario: longitudinal volumetry. Specifically, we first used SynthSR and NiftyReg to super-resolve and register the low-field scans across timepoints. The resulting images were then segmented with SynthSeg^45^, and the corresponding segmentations were warped to the opposite timepoint using the estimated deformation fields. For each timepoint, we fused the native and propagated segmentations using a label fusion method that applies Gaussian weighting based on local intensity differences^46^, following the approach successfully employed in our previous work^47^. The resulting soft segmentations were then used to compute final volumetric estimates for all structures segmented by SynthSeg.

Figure 5 shows spaghetti plots showing the longitudinal volume trajectories of the hippocampus, ventricles, and whole brain. In the previous experiments, the registration used to correct geometric distortion also largely mitigated the temporal gaps between the low- and high-field acquisitions. In contrast, in the present setup, these temporal gaps prevent direct comparison with the high-field references. Nevertheless, the plots demonstrate that low-field MRI, when combined with longitudinal segmentation methods, can capture expected aging-related trends such as ventricular expansion and atrophy of the hippocampus and whole brain, particularly at 3 mm isotropic resolution. A few trajectories deviate from the expected direction, but these typically correspond to subjects with shorter inter-scan intervals, where measurement variability has a proportionally greater effect.

**Figure 5.**
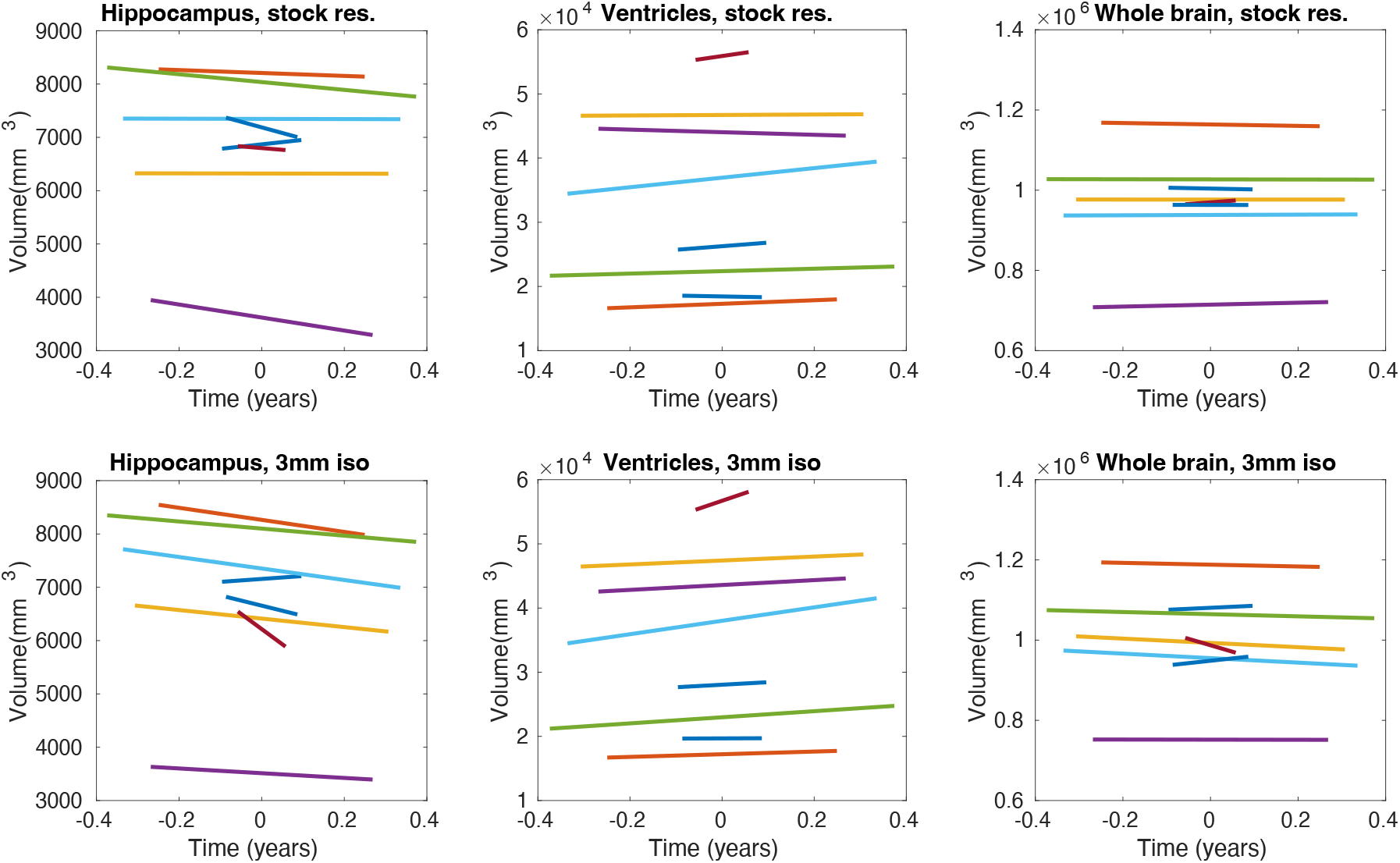
Spaghetti plots for longitudinal trajectories of volumes for the bilateral hippocampus, lateral ventricles, and whole brain of eight subjects, using T1 low-field MRI scans acquired at stock (1.6 × 1.6 × 5.0 mm axial) and 3 mm isotropic resolution. Each color represents a different subject.

## Discussion

This paper experimentally tests whether existing registration methods are accurate enough to analyze low-field brain MRI scans acquired with a portable device. We consider three different representation methods: direct registration via classical optimization, direct registration via neural networks, and indirect registration via synthesis of 1 mm isotropic images followed by classical registration. Our results demonstrate that existing registration methods can effectively handle the limited resolution of low-field MRI. When the resolution reduction is simulated by downsampling high-field data, all approaches achieve high accuracy and smooth deformation fields. This indicates that image resolution alone is not the limiting factor for registration performance in low-field imaging. In contrast, real-world low-field scans present a much greater challenge. The performance degradation observed in these data reflects the presence of acquisition artifacts such as strong intensity inhomogeneity, noise, and particularly geometric distortion, which are not captured by simple downsampling. Among the evaluated methods, only the synthesis-based approach (SynthSR combined with NiftyReg) provided results that were consistent and usable for most subjects. SynthMorph, while performing well on simulated low-resolution data, was not trained on images with the noise characteristics or distortions typical of low-field MRI, which may explain its reduced performance in this setting.

These challenges become particularly important in longitudinal analyses, where accurate registration across time points is essential to detect subtle anatomical changes. Our experiments show that performance decreases with real low-field scans, especially at anisotropic resolutions. For longitudinal studies, we therefore recommend using isotropic acquisitions whenever possible, combined with synthesis-based registration methods and established longitudinal analysis frameworks, such as the one used here or complementary techniques like linear mixed-effects modeling.

Future work will focus on three main directions. First, improved characterization of geometric distortion through expanded phantom studies will be essential for quantifying and modeling the underlying low-field artifacts. Second, new computational methods are needed to explicitly account for these distortions, ideally by separating scanner-induced geometric warping from true anatomical variation. One promising avenue is to retrain SynthMorph or related neural registration frameworks using simulations that incorporate biological variability together with low-field–specific signal and artifact models. Importantly, these computational advances are complementary to ongoing hardware and calibration efforts aimed at reducing distortion at the acquisition stage; together, they could substantially improve the fidelity of downstream registration. Third, broader evaluations on larger and more diverse datasets will help establish the practical limits, robustness, and variability of registration performance under real-world low-field conditions.

Ultimately, as the demand for frequent brain imaging grows (e.g., for monitoring aging, tracking neurodegenerative diseases, or assessing responses to treatments such as Alzheimer’s therapeutics), the ability to perform reliable registration on low-cost, portable MRI scanners becomes increasingly important. Achieving robust registration at low field is not only a technical milestone but a foundational requirement for enabling global-scale, quantitative neuroimaging studies that extend far beyond traditional hospital settings.

## Materials and Methods

### MRI datasets

#### ADNI3

We used near 1 mm isotropic high-field (3 T) T1 and FLAIR scans from 80 randomly selected subjects from the third phase of the Alzheimer’s Disease Neuroimaging Initiative (ADNI, adni.loni.usc.edu): 43 males vs 37 females; average age 78.7 ±7.0 years; and 41 elderly controls vs 39 subjects with some degree of cognitive impairment. Segmentations for 31 brains regions were automatically obtained using the robust version^48^ of our publicly available SynthSeg tool^45^, which is agnostic to MRI contrast and resolution, and thus compatible with both T1 and FLAIR scans. The ADNI was launched in 2003 as a public-private partnership, led by Principal Investigator Michael W. Weiner, MD. The primary goal of ADNI has been to test whether serial MRI, positron emission tomography, other biological markers, and clinical and neuropsychological assessment can be combined to measure the progression of mild cognitive impairment, and early Alzheimer’s disease. For up-to-date information, see www.adni-info.org.

#### Study Design for Low-Field Datasets

Two low-field patient datasets were prospectively acquired at the Massachusetts General Hospital (Low-Field Outpatient Dataset) and Yale New Haven Hospital (Longitudinal Dataset) between February 2023 and May 2025. Exclusion criteria were electrically active implants or a body weight exceeding 400 lbs. Participants were enrolled under Mass General Brigham or Yale New Haven Hospital Institutional Review Board approved protocol with informed consent obtained prospectively. All experiments were performed in accordance with relevant guidelines and regulations.

#### Low-Field Outpatient Dataset

This dataset comprises low-field MRI scans from 40 subjects (average age 65.3 ± 15.1 years, 21 female vs 19 male) acquired with a 64 mT Swoop^®^ scanner version Mk1.9 (Hyperfine, Inc), and paired high-field scans acquired at 1.5-3 T for clinical care (Siemens, GE and Philips). Patients were recruited between February 2023 and December 2024 and scanned in an outpatient neurology clinic at Massachusetts General Hospital, with images acquired following routine appointments. Eligible participants included those with physician-diagnosed neurodegenerative disease, comprising mild cognitive impairment or dementia due to Alzheimer’s disease, multiple sclerosis, and Parkinson’s disease. The low-field scans were acquired with T1 and FLAIR contrast, at both “stock” (1.6 × 1.6 × 5 mm axial) and 3 mm isotropic resolution. The high-field sessions included 1 mm isotropic T1 and FLAIR scans. As for ADNI3, automated segmentations were obtained from the high-field scans using SynthSeg. These segmentations were propagated to the low-field scans via registration, using NiftyReg with LNCC and a 20 mm B-Spline control point spacing (as per the results of the phantom experiment).

#### Longitudinal Dataset

This dataset comprises low-field MRI scans from 8 subjects (average age 67.1±13.7 years, 5 female vs 3 male) acquired with a 64 mT Swoop^®^ scanner version Mk1.9 (Hyperfine, Inc), as well as paired high-field scans of the same subjects acquired on different 3 T Siemens Magnetom scanners (including Verio, Vida, and Skyra). Patients were recruited between May 2024 and May 2025 and scanned at a tertiary care memory clinic at Yale New Haven Hospital. Eligible participants included individuals with physician-diagnosed mild cognitive impairment or Alzheimer’s disease. Diagnosis was determined by neurologists based on clinical evaluation and standard diagnostic criteria. The low-field scans were acquired with T1 contrast at both “stock” (1.6 × 1.6 × 5 mm axial) and 3 mm isotropic resolution. The high-field sessions included only FLAIR scans, acquired axially with sub-mm in-plane resolution, and slice spacings between 1 and 5 mm (average: 3.7 ± 1.3 mm). As for the other datasets, automated segmentations were obtained from the high-field scans using SynthSeg, and propagated to the low-field scans via registration with a highly constrained nonlinear model (NiftyReg with LNCC and a 20 mm B-Spline control point spacing). We note that, since the high-field scans are not isotropic, they were super-resolved to 1 mm isotropic with SynthSR prior to registation when generating the HF-NiftyReg reference.

#### Phantom

This dataset utilized a cylindrical propylene glycol phantom provided by the manufacturer (Hyperfine, Inc). The phantom was first scanned at high field using a T1, a T2, and a FLAIR sequence, all at 1 mm isotropic resolution on a 3 T Siemens Prisma fit scanner with a 64-channel head-coil.

The phantom was also scanned five times at low field, with T1, T2, and FLAIR contrast, at stock resolution (1.6 × 1.6 × 5 mm axial). These five sessions relied on five different combinations of hardware and software versions of the Swoop^®^ scanner: Mk1.6/rc8.5.0, Mk1.6/8.3.0, Mk1.6/8.4.0, Mk1.9/rc8.6.0, and Mk1.9/rc8.6.0.

### Registration methods

We considered three open-source, state-of-the-art methods, which cover three representative families of algorithms: classical optimization (NiftyReg), learning-based registration (SynthMorph), and synthesis-based registration (SynthSR+NiftyReg).

- **NiftyReg**^16^ (version 1.5.77): this is a classical method based on numerical optimization. We used the diffeomorphic version with LNCC (*σ* =7 mm) within modalities and normalized mutual information (64 bins) across modalities. When registering high-field scans, we used a pyramid with three levels and 5 mm B-Spline control spacing at the finest level. With the low-field scans, we used the same control point spacing but only two levels. Prior to registration, scans were skull-stripped with SynthSeg.
- **SynthMorph**^41^ (version distributed with FreeSurfer 8): this is a deep learning method based on amortized optimization with a neural network. SynthMorph is trained to register brain scans using synthetic, domain-randomized data (similar to SynthSeg). This makes the method usable “out of the box” (i.e., without finetuning or adapting), since it is agnostic to changes in resolution and contrast.
- **SynthSR+NiftyReg** (SynthSR distributed with FreeSurfer 8 and NiftyReg 1.5.77, as above): SynthSR^42^ relies on the same domain randomization methods as SynthSeg and SynthMorph, and estimates (also “out of the box”) a 1 mm isotropic T1 volume from a brain MRI of any resolution and contrast – which can be subsequently registered with NiftyReg using the intra-modality configuration described above. Previous studies have shown that the synthesis error is largely offset by the gain in registration accuracy, particularly when performing nonlinear registration across different resolutions or MRI contrasts^44^.

### Metrics

Overlap of segmentations is measured with Dice coefficients:

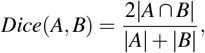

where | · | denotes cardinality. This metric ranges from zero (no overlap) to one (perfect overlap).

The regularity of deformation fields is assessed with membrane energies, which measure the level of stretching of the deformation. We compute the membrane energy over a domain Ω equal to the brain region, and normalize it over its area for comparison across subjects:

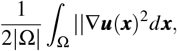

where ***u*** is the deformation field and ***x*** represents the spatial coordinates. In the case of the phantom, since we are scanning the same object twice, we instead use log-Jacobians: since we expect the transformation to preserve volume, we measure deviations from the expected unit Jacobian determinant of ***u***(***x***).

## Funding Declaration

This work was funded by NIH grants R01-AG070988, R01-EB031114, UM1-MH130981, RF1-AG080371, R21-NS138995, R01-EB031114, and R21-NS145048.

## Acknowledgements

The data collection and sharing of the ADNI data was funded by the Alzheimer’s Disease Neuroimaging Initiative (National Institutes of Health Grant U01-AG024904) and DOD ADNI (Department of Defense award number W81XWH-12-2-0012). ADNI is funded by the National Institute on Aging, the National Institute of Biomedical Imaging and Bioengineering, and through generous contributions from the following: AbbVie, Alzheimer’s Association; Alzheimer’s Drug Discovery Foundation; Araclon Biotech; BioClinica, Inc.; Biogen; Bristol-Myers Squibb Company; CereSpir, Inc.; Cogstate; Eisai Inc.; Elan Pharmaceuticals, Inc.; Eli Lilly and Company; EuroImmun; F. Hoffmann-La Roche Ltd and its affiliated company Genentech, Inc.; Fujirebio; GE Healthcare; IXICO Ltd.; Janssen Alzheimer Immunotherapy Research & Development, LLC.; Johnson & Johnson Pharmaceutical Research & Development LLC.; Lumosity; Lundbeck; Merck & Co., Inc.; Meso Scale Diagnostics, LLC.; NeuroRx Research; Neurotrack Technologies; Novartis Pharmaceuticals Corporation; Pfizer Inc.; Piramal Imaging; Servier; Takeda Pharmaceutical Company; and Transition Therapeutics. The Canadian Institutes of Health Research is providing funds to support ADNI clinical sites in Canada. Private sector contributions are facilitated by the Foundation for the National Institutes of Health (www.fnih.org). The grantee organization is the Northern California Institute for Research and Education, and the study is coordinated by the Alzheimer’s Therapeutic Research Institute at the University of Southern California. ADNI data are disseminated by the Laboratory for Neuro Imaging at the University of Southern California.

## Author contributions statement

1. **Conceptualization**: JEI, IPJ, KG, LT, MSR, KNS, AdH, WTK, ASA.
2. **Data curation**: JEI, IPJ, JWR, DZ, MO, ADF, AD, ASA.
3. **Formal analysis**: JEI, ASA.
4. **Funding acquisition**: JEI, MSR, KNS, AdH, WTK, ASA.
5. **Investigation**: IPJ, MO, ADF, AD.
6. **Methodology**: JEI, LT, KG, ASA.
7. **Project administration**: JEI, ASA.
8. **Resources**: MSR, KNS, WTK.
9. **Writing – original draft**: JEI.
10. **Writing – review and editing**: All authors.

## Additional information

### Competing interests

MSR has an interest in Hyperfine, Inc. The Yale-affiliated authors acknowledge funding from Hyperfine and Genentech to support their portable MRI program.

## Supplementary Figures

**Figure S1.**
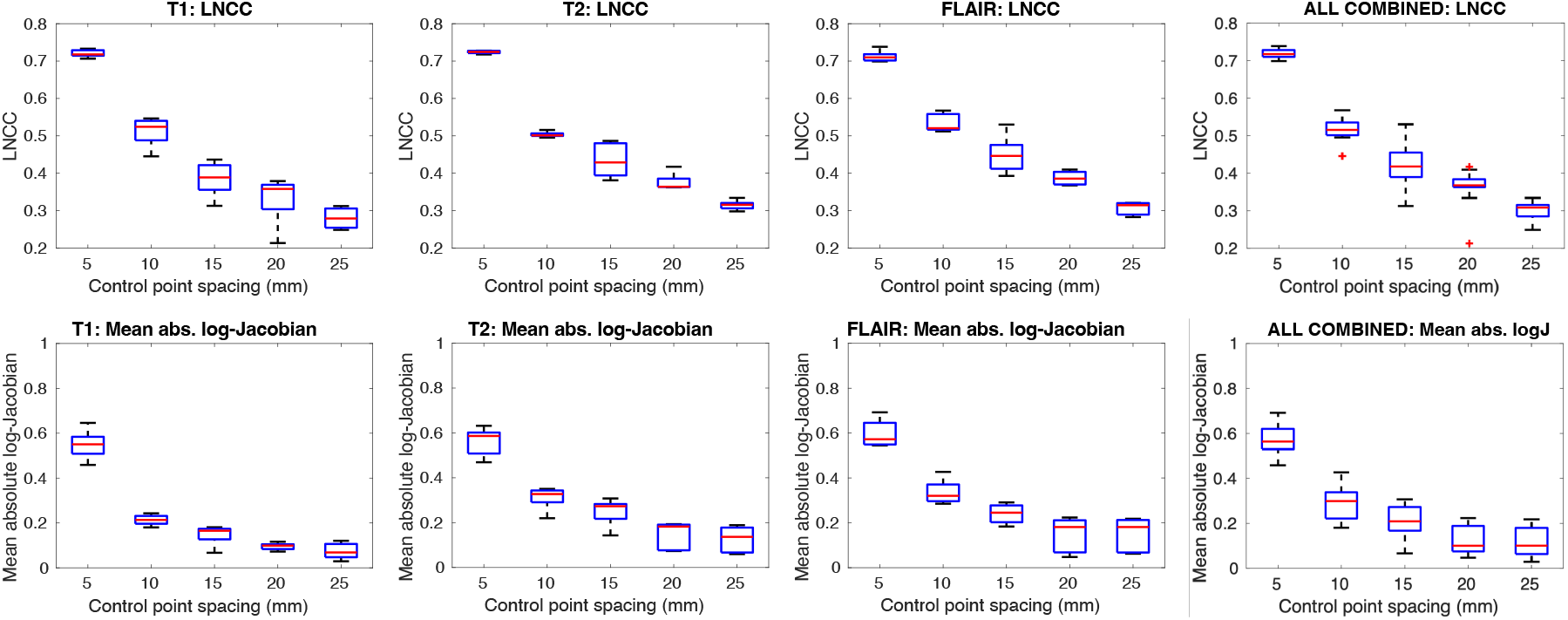
Box plots for LNCC and mean absolute log-Jacobian for phantom registration from high to low field, over five different low-field sessions (details in Methods), for five different control point spacings of the B-Spline transform in NiftyReg. We show plots for T1, T2, and FLAIR scans, as well as for all pulse sequences combined. The boxes show medians (in red), interquartile ranges (IQR, in blue), whiskers (±1.5 × IQR, in black), and outliers (red crosses). We selected 20 mm as a compromise, since smaller spacings led to rapidly increasing irregularity, and the 20 mm transform sufficed to achieve accurate visual alignment (Figure 2d), i.e., higher LNCC values seem to indicate overfitting.

## Notes

http://www.adni-info.org/

## References

1. Fitzpatrick, J. M., West, J. B. & Maurer, C. R. Predicting error in rigid-body point-based registration. IEEE transactions on medical imaging 17, 694–702 (2002).

2. Pluim, J. P., Maintz, J. A. & Viergever, M. A. Mutual-information-based registration of medical images: a survey. IEEE transactions on medical imaging 22, 986–1004 (2003).

3. Holland, D., Dale, A. M., Initiative, A. D. N. et al. Nonlinear registration of longitudinal images and measurement of change in regions of interest. Med. image analysis 15, 489–497 (2011).

4. James, A. P. & Dasarathy, B. V. Medical image fusion: A survey of the state of the art. Inf. fusion 19, 4–19 (2014).

5. Iglesias, J. E. & Sabuncu, M. R. Multi-atlas segmentation of biomedical images: a survey. Med. image analysis 24, 205–219 (2015).

6. Chen, M., Tustison, N. J., Jena, R. & Gee, J. C. Image registration: Fundamentals and recent advances based on deep learning. Mach. Learn. for Brain Disord. 435–458 (2023).

7. Fonov, V. S., Evans, A. C., McKinstry, R. C., Almli, C. R. & Collins, D. Unbiased nonlinear average age-appropriate brain templates from birth to adulthood. NeuroImage 47, S102 (2009).

8. Ashburner, J. & Friston, K. J. Voxel-based morphometry—the methods. Neuroimage 11, 805–821 (2000).

9. Klein, A. et al. Evaluation of 14 nonlinear deformation algorithms applied to human brain MRI registration. Neuroimage 46, 786–802 (2009).

10. Evans, A. C., Janke, A. L., Collins, D. L. & Baillet, S. Brain templates and atlases. Neuroimage 62, 911–922 (2012).

11. Thirion, J.-P. Image matching as a diffusion process: an analogy with Maxwell’s demons. Med. image analysis 2, 243–260 (1998).

12. Rueckert, D. et al. Nonrigid registration using free-form deformations: application to breast MR images. IEEE transactions on medical imaging 18, 712–721 (2002).

13. Avants, B. B., Epstein, C. L., Grossman, M. & Gee, J. C. Symmetric diffeomorphic image registration with cross-correlation: evaluating automated labeling of elderly and neurodegenerative brain. Med. image analysis 12, 26–41 (2008).

14. Viergever, M. A. et al. A survey of medical image registration–under review (2016).

15. Sotiras, A., Davatzikos, C. & Paragios, N. Deformable medical image registration: A survey. IEEE transactions on medical imaging 32, 1153–1190 (2013).

16. Modat, M. et al. Fast free-form deformation using graphics processing units. Comput. methods programs biomedicine 98, 278–284 (2010).

17. Klein, S., Staring, M., Murphy, K., Viergever, M. A. & Pluim, J. P. Elastix: a toolbox for intensity-based medical image registration. IEEE transactions on medical imaging 29, 196–205 (2009).

18. Andersson, J., Jenkinson, M., Smith, S. et al. Non-linear registration, aka spatial normalisation FMRIB technical report TR07JA2. FMRIB Analysis Group Univ. Oxf. 2, e21 (2007).

19. Ashburner, J. A fast diffeomorphic image registration algorithm. Neuroimage 38, 95–113 (2007).

20. Alfaro-Almagro, F. et al. Image processing and quality control for the first 10,000 brain imaging datasets from UK Biobank. Neuroimage 166, 400–424 (2018).

21. LeCun, Y., Bottou, L., Bengio, Y. & Haffner, P. Gradient-based learning applied to document recognition. Proc. IEEE 86, 2278–2324 (2002).

22. Vaswani, A. et al. Attention is all you need. Adv. neural information processing systems 30 (2017).

23. Sokooti, H. et al. Nonrigid image registration using multi-scale 3D convolutional neural networks. In International conference on medical image computing and computer-assisted intervention, 232–239 (Springer, 2017).

24. Yang, X., Kwitt, R., Styner, M. & Niethammer, M. Quicksilver: Fast predictive image registration–a deep learning approach. NeuroImage 158, 378–396 (2017).

25. Balakrishnan, G., Zhao, A., Sabuncu, M. R., Guttag, J. & Dalca, A. V. Voxelmorph: a learning framework for deformable medical image registration. IEEE transactions on medical imaging 38, 1788–1800 (2019).

26. De Vos, B. D. et al. A deep learning framework for unsupervised affine and deformable image registration. Med. image analysis 52, 128–143 (2019).

27. Fu, Y. et al. Deep learning in medical image registration: a review. Phys. Medicine & Biol. 65, 20TR01 (2020).

28. Haskins, G., Kruger, U. & Yan, P. Deep learning in medical image registration: a survey. Mach. Vis. Appl. 31, 8 (2020).

29. Dalca, A. V., Balakrishnan, G., Guttag, J. & Sabuncu, M. R. Unsupervised learning of probabilistic diffeomorphic registration for images and surfaces. Med. image analysis 57, 226–236 (2019).

30. Jiang, Z., Yin, F.-F., Ge, Y. & Ren, L. A multi-scale framework with unsupervised joint training of convolutional neural networks for pulmonary deformable image registration. Phys. Medicine & Biol. 65, 015011 (2020).

31. Chen, J. et al. Transmorph: Transformer for unsupervised medical image registration. Med. image analysis 82, 102615 (2022).

32. Kim, B., Han, I. & Ye, J. C. Diffusemorph: Unsupervised deformable image registration using diffusion model. In European conference on computer vision, 347–364 (Springer, 2022).

33. Ho, J., Jain, A. & Abbeel, P. Denoising diffusion probabilistic models. Adv. neural information processing systems 33, 6840–6851 (2020).

34. Tian, L. et al. uniGradICON: A foundation model for medical image registration. In International Conference on Medical Image Computing and Computer-Assisted Intervention, 749–760 (Springer, 2024).

35. Turpin, J. et al. Portable magnetic resonance imaging for ICU patients. Critical Care Explor. 2, e0306 (2020).

36. Yuen, M. M. et al. Portable, low-field magnetic resonance imaging enables highly accessible and dynamic bedside evaluation of ischemic stroke. Sci. advances 8, eabm3952 (2022).

37. Kimberly, W. T. et al. Brain imaging with portable low-field MRI. Nat. reviews bioengineering 1, 617–630 (2023).

38. Srinivas, S. A. et al. External dynamic interference estimation and removal (EDITER) for low field MRI. Magn. resonance medicine 87, 614–628 (2022).

39. Sorby-Adams, A. J. et al. Portable, low-field magnetic resonance imaging for evaluation of Alzheimer’s disease. Nat. Commun. 15, 10488 (2024).

40. Wells III, W. M., Viola, P., Atsumi, H., Nakajima, S. & Kikinis, R. Multi-modal volume registration by maximization of mutual information. Med. image analysis 1, 35–51 (1996).

41. Hoffmann, M. et al. SynthMorph: learning contrast-invariant registration without acquired images. IEEE transactions on medical imaging 41, 543–558 (2021).

42. Iglesias, J. E. et al. SynthSR: A public AI tool to turn heterogeneous clinical brain scans into high-resolution T1-weighted images for 3D morphometry. Sci. advances 9, eadd3607 (2023).

43. Szeliski, R. Computer vision: algorithms and applications (Springer Nature, 2022).

44. Iglesias, J. E. et al. Is synthesizing MRI contrast useful for inter-modality analysis? In International Conference on Medical Image Computing and Computer-Assisted Intervention, 631–638 (Springer, 2013).

45. Billot, B. et al. SynthSeg: Segmentation of brain MRI scans of any contrast and resolution without retraining. Med. image analysis 86, 102789 (2023).

46. Sabuncu, M. R., Yeo, B. T., Van Leemput, K., Fischl, B. & Golland, P. A generative model for image segmentation based on label fusion. IEEE transactions on medical imaging 29, 1714–1729 (2010).

47. Casamitjana, A. et al. USLR: An open-source tool for unbiased and smooth longitudinal registration of brain MRI. Med. Image Analysis 103662 (2025).

48. Billot, B. et al. Robust machine learning segmentation for large-scale analysis of heterogeneous clinical brain MRI datasets. Proc. Natl. Acad. Sci. 120, e2216399120 (2023).

